# Interaction of human CRX and NRL in live HEK293T cells measured using fluorescence resonance energy transfer (FRET)

**DOI:** 10.1101/2021.10.19.465002

**Authors:** Xinming Zhuo, Barry E. Knox

**Affiliations:** Departments of Ophthalmology & Visual Sciences and Biochemistry & Molecular Biology, Center for Vision Research, SUNY Upstate Medical University, Syracuse, NY 13210, USA

## Abstract

CRX and NRL are retina-specific transcription factors that control rod photoreceptor differentiation and synergistically activate rod phototransduction gene expression. Previous experiments showed they interact *in vitro* and in yeast two-hybrid assays. Here, we examined CRX-NRL interaction in live HEK293T cells using two fluorescence resonance energy transfer (FRET) approaches: confocal microscopy and flow cytometry (FC-FRET). FC-FRET can provide measurements from many cells having wide donor-acceptor expression ranges. FRET efficiencies were calibrated with a series of donor (EGFP)-acceptor (mCherry) fusion proteins separated with linkers between 6-45 amino acids. CRX and NRL were fused at either terminus with EGFP or mCherry to create fluorescent proteins, and all combinations were tested in transiently transfected cells. FRET signals between CRX or NRL homo-pairs were highest with both fluorophores fused to the DNA binding domains (DBD), lower with both fused to the activation domains (AD), and not significant when fused on opposite termini. NRL had stronger FRET signals than CRX. A significant FRET signal between CRX and NRL hetero-pairs was detected when donor was fused to the CRX DNA binding domain and the acceptor fused to the NRL activation domain. FRET signals increased with CRX or NRL expression levels at a rate much higher than expected for collisional FRET alone. Together, our results show the formation of CRX-NRL complexes in live HEK293T cells that are close enough for FRET.

## Introduction

Vertebrate photoreceptors express a large array of genes [1] specifically related to phototransduction [2] and their unique cellular structures [3, 4], such as the outer segment [5, 6]. CRX [7-9], an OTX-like protein that is a member of the *paired* homeodomain family, and NRL [10, 11], a basic leucine zipper (bZIP) protein that is a member of the large Maf family, are key retinal transcription factors essential for photoreceptor function. Together, they regulate rod photoreceptor differentiation and gene expression [12, 13], are involved in the *in vitro* differentiation of stem cells into photoreceptors [14-17] and are implicated in human retinal diseases [18-22]. Moreover, CRX and NRL are expressed in medulloblastoma cells, where they activate photoreceptor genes and contribute to tumor maintenance [23]. In addition to a direct role in causing retinal disease via alterations of their protein sequence, they also play an indirect role by regulating genes that cause inherited retinal degenerative diseases. For example, there are more than 90 genes linked to one or more of six commonly occurring retinal diseases [24]. Many of these genes are directly regulated by CRX or NRL [25] or have putative cis-regulatory DNA binding sites close to their transcription initiation sites [26]. CRX and NRL together regulate transcription initiation of numerous genes [26, 27] directly by binding to *cis*-regulatory elements in promoter regions [26-29], indirectly through chromatin modification [30-32], and by interacting with or regulating other transcription factors [10, 25].

A thorough understanding of CRX and NRL structure and function is essential, not only for establishing the mechanistic basis of photoreceptor gene expression, but for developing new treatments for human disease. Genome-wide analysis has identified a consensus CRX [26, 33] and NRL [34, 35] cis-regulatory sequences that cluster in or near rod photoreceptor genes, suggesting that CRX and NRL together regulate them [26, 33]. The localization of CRX-NRL sites in proximal promoter regions reinforces functional and biochemical experiments that demonstrate action by CRX and NRL to increase transcription [7, 36-38]. In transiently transfected cultured cell lines, CRX and NRL can individually activate transcription from rhodopsin and other photoreceptor-specific promoters, but together they do so synergistically [7, 36, 37]. CRX and NRL can bind to each other *in vitro* in the absence of DNA and can interact in yeast cells as inferred from two-hybrid studies [36]. Although the NRL bZIP domain and the CRX homeodomain have roles in CRX-NRL interaction *in vitro*, interaction appears to involve other regions of both proteins as well [36]. Little is known about the underlying structural interface(s) that mediate complex formation, the structural basis for their transcriptional activity, protein-DNA or protein-protein interactions. The importance in understanding the structure-function relationships that result in CRX-NRL cooperative transcriptional activity is highlighted by the fact that mutations in CRX or NRL that reduce synergistic transactivation in cell transfection assays are linked to human retinopathies [20].

In this report, we describe the characterization of the interactions between CRX and NRL in cultured mammalian cells. We utilized transiently transfected HEK293 cells because they are readily transfected and have been used in numerous studies to functionally characterize CRX-NRL [7, 36, 37]. Most importantly, CRX and NRL are able to activate co-transfected but not endogenous phototransduction gene promoters. Previous studies used bioluminescence resonance energy transfer (BRET), a technique to study the interactions of Nr2e3 with itself, CRX and NRL in pooled populations of HEK293 cells [39, 40]. The ability to directly image live HEK293 cells and to readily sort them with minimal manipulations offer an advantage for initial investigations of FRET between these two transcription factors. This approach is suitable for detecting protein-protein interactions, without distinguishing whether the interactions occur bound to DNA/chromatin or free in the nucleus. To measure FRET in living cells [41], we used either an improved FC-FRET approach, described here, or confocal microscopy FRET, CM-FRET [42].

CM-FRET offers subcellular spatial resolution and the potential to observe movements of FRET partners by photobleaching methods (reviewed in [43, 44]). However, data collection with CM-FRET can be limited by both the number of cells that can be processed and biases in cell selection. Previously, flow cytometry has been used for analyzing FRET in populations of cells [45-53], including the interaction of transcription factors [51, 54]. We adapted one FC-FRET method [51] in order to measure sensitized emission derived from donor-acceptor pairs and calibrated it using mCherry-EGFP (mG) fusion proteins separated by different length linkers. Using a combination of microscopy and flow cytometry, we characterized the interactions in living cells of CRX and NRL fused to mCherry or EGFP and show interactions between these two transcription factors that are close enough for FRET.

## Results

### Measurement of FRET by flow cytometry

Transiently transfected HEK293T cells have served as a convenient and rapid model system for the characterization of retinal transcription factors (e.g. [37]). We adapted an FC-FRET approach to measure apparent FRET efficiencies (N_FRET_) determined by sensitized emission. For these studies, EGFP served as the donor and mCherry as the acceptor. Since our goal was to examine FRET between transcription factors, all protein constructs had a nuclear localization signal added to the N-terminus to direct expression exclusively to the nucleus. For each FC-FRET experiment (Figure S1), four control groups of cells were transfected with the following expression constructs: mock (empty pcDNA3.1), mCherry alone, EGFP alone, and unlinked mCherry plus EGFP (mCh+eG). Previous approaches estimated FRET by counting the number of cells that cross a threshold level of corrected F^DA^ intensity (Figure S2). In order to quantify FRET signals, we employed a well-established method (three cube, [55]) to determine N_FRET_ on a cell-by-cell basis. Subsequently, the population averages and partitions based upon donor-acceptor intensity levels were used. The four control groups were used to estimate expression levels (Figure S3) and background–bleed-through fluorescence (crosstalk) in the FRET channel (Figure S4). To calibrate actual FRET signals (Figures 1, S4, and S5), we used donor-acceptor fusion proteins (mG) which undergo intramolecular FRET when EGFP is excited. The fusion was accomplished through linkers (Supplemental Table 1) containing an α-helix-forming peptide, EAAAK [56] repeated 2 to 7 times, and flanked on each side by a proline residue to terminate the α-helical region. For both unlinked and mG constructs, fluorescence was observed uniformly in the nucleoplasm (Figure S6). There was an enrichment in the nucleolus compared to the nucleoplasm, with a mean ratio of ∼2 for both donor and acceptor fluorescence (*data not shown*). This is consistent with the behaviour of NLS which mediate RNA binding and nucleolar localization of fluorescent proteins [57].

**Figure 1.**
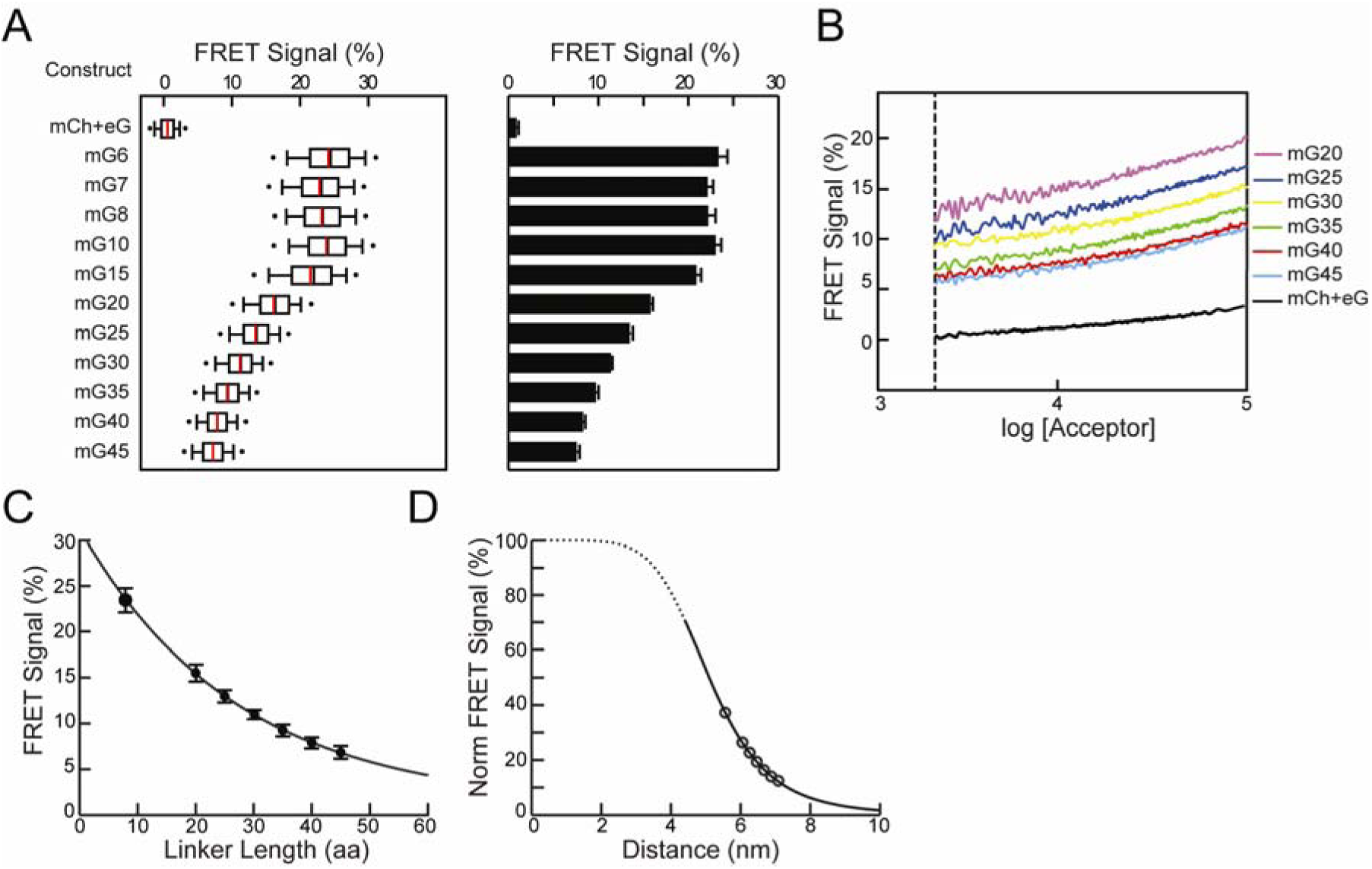
Flow cytometry FRET. A) Comparison of FRET signals (N_FRET_) between mCherry and EGFP fusion constructs (mGx) that contain linkers of different length (x = number of amino acids) in a single flow cytometry run (*left panel*) or averaged over multiple experiments (*right panel*). For the single experiment, results are shown as box plot, with mean (*red lines*) and median (*black lines*) values indicated, (number of cells: 4385-10215). For averaged experiments, mean values with SEM of the FRET signals are shown in a bar graph, (number of flow runs: 3-5). B) The FRET signals from cells expressing mG20-45 from a single flow cytometry run. The distribution range of cell acceptor fluorescence intensities was divided into intervals with 100 cells in each bin. In the intensity plots, moving averages of N_FRET_ were calculated and are plotted versus the acceptor intensity in each bin. Error bars are suppressed for clarity. C) Mean N_FRET_ with SEM from the averaged experiments (A) are plotted as a function of linker length for mG8 and mG20-45. The line is a fit (R^2^= 0.998) to the Förster equation modified to use linker length with the parameters k_1_=0.27, k_2_=5.56. (D) Normalized FRET signals from C plotted as a function of predicted distance fit to the FRET equation (*solid black line*) with the same parameters in C. Dashed line indicates the inaccessible distance between mCherry and EGFP due to steric volume exclusion between the two fluorescent proteins.

FC-FRET produces sufficient cell numbers to restrict analysis to those that optimize the FRET signal to noise ratio (*e*.*g*., Figure 1). To control for fluorescent protein expression levels which ranged over three orders of magnitude (<10^−7^ to ∼2 ×10^−4^ M, Figure S4), low expressing cells (with donor/acceptor fluorescence <∼3 × 10^3^ FU, Figures S4, S5) were eliminated from analysis (Figure S5). Only cells with an expression ratio for mCherry:EGFP between 0.1-10, the range over which the FRET efficiency is most stable [58], were included. A criterion F^DA^ level (fluorescence intensity in the acceptor channel when excited with the donor laser) was set so that no cells that expressed either mCherry or EGFP alone reached this F^DA^ intensity (Figure S5, similar to a previous report [45]). Cells that meet the above criteria F^DA^ intensity were termed FRET-positive cells. With this optimization, ∼80.0% mG fusion construct expressing cells and ∼4-5% of mCh+eG expressing cells were classified as FRET-positive in typical experiments (Figures S4 and S5) and were used to calculate N_FRET_. In a typical experiment, we observed a mean N_FRET_ for mG10 expressing cells, expected to have a high FRET efficiency, that ranged from 21-24% while mCh+eG cells, expected to exhibit background FRET efficiency, was less than 1.5% (Figures 1A and S5). This represents an order of magnitude range for comparison of N_FRET_ in transfected HEK293T cells.

In addition to the intrinsic FRET that depends on the close proximity of donor-acceptor fluorophores, stochastic or collisional FRET arises from transient interactions between donor and acceptor [59]. Stochastic FRET is expected to linearly depend upon the concentration of freely diffusing donors and acceptors [59-61]. To estimate the contribution of stochastic FRET to the signal measured by flow cytometry, we examined cells expressing mG fusion proteins with a wide range of acceptor and donor fluorescence levels (Figure S6). This is readily accomplished since the FRET signals are collected over the entire range of mG expression during flow cytometry (Figure S7). In cells expressing mCh+eG, N_FRET_ modestly increased as either acceptor (Figures 1B and S7A) or donor (Figure S7B) fluorescence levels increased. The dependence of N_FRET_ on acceptor concentration fit the stochastic FRET equation ([59-61], Figure S8), indicating that the mCh+eG samples gave an accurate measure of the stochastic FRET component. We also compared collisional FRET in cells expressing nuclear-localized mCh+eG with those expressing cytoplasmic mCh+eG and found the dependence on expression level was indistinguishable in the two cellular compartments (Figure S9), further supporting the identification of the mCh+eG signal with stochastic FRET. To characterize intrinsic FRET in the following experiments, we compared a population measure of FRET encompassing a range of expression levels (N_FRET_) and the FRET efficiency dependence on fluorophore concentration.

### Measurement of distance by FC-FRET

To quantify FRET efficiencies as a function of donor-acceptor distance, we analysed N_FRET_ from cells expressing mG fusion proteins with different linker lengths (Figure 1A, Figure S6). The linker design incorporated a rigid alpha helix [56] that has been studied by X-ray analysis and shown to influence FRET efficiency in a fusion protein between BFP and GFP *in vitro* [62-64]. The two proline residues incorporated in the mG design should isolate the alpha helical segments proteins to reduce influences of relative orientation of the two fluorophores. Fusion proteins with linkers less than 15 amino acids all exhibited similar N_FRET_ either in single or when averaging multiple FC-FRET experiments (mG6-8, one way ANOVA: α=0.05, F=1.54, p=0.29; mG10 and mG15, t-test: mG-8 vs. mG10, p=0.227; mG-8 vs. mG15, p=0.116). N_FRET_ was very sensitive to lengths longer than 15 amino acids, with a steady reduction as linkers lengthened in individual (Figure 1A, *left*) or combined (Figure 1A, *right*) FC-FRET experiments (one way ANOVA, Holm-Sidak, pairwise comparison, all pairs have p<0.05).

All mG fusion proteins exhibited an increase in FRET efficiency as acceptor (Figures 1B and S7A) or donor (Figure S7B) fluorescence intensities increased, but the vertical offsets and slopes varied. The mG series of fusion proteins had parallel curves that differed in the offset at all fluorescence levels (Figures 1B and S7). This offset depended upon the linker length and represented the intrinsic FRET from the donor-acceptor pairs. There was a gradual increase in the slope at higher expression levels which was similar for all mG fusion proteins and was greater than that for mCh+eG. The cause of the increasing slope at higher tethered mCh+eG concentrations compared to untethered fluorophores is not clear; it may be due to multiple donor-acceptor interactions, either from concentration dependent association of fluorophores or collisions between tethered fluorophores. The effect of donor-acceptor expression level on FRET signals highlights the importance of comparing donor-acceptor pairs in the same concentration range to clearly distinguish the intrinsic and stochastic FRET signals in transfected cells [59-61].

To estimate the dependence of FRET efficiency on distance, R, between donor and acceptor, we fit N_FRET_ to the Förster equation (Figure 1C):

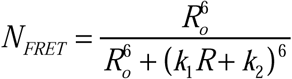 where R_o_ is Förster distance (5 nm for EGFP and 4.7-5.2 nm for mCherry [65-67]), k_1_ an orientation factor between the fluorescent proteins, and k_2_ is the minimal distance between two fluorophores determined by steric exclusion. The data was well fit to this equation using the predicted lengths of the alpha helical linkers [62-64]. These results show that FC-FRET can quantitatively measure small differences in distance between donors and acceptors in living cells.

### Comparison of flow cytometry and confocal microscopy FRET

We used both confocal microscopy and flow cytometry FRET with mG10 (Figure 2) to quantitatively compare: i) FRET signal/efficiency; ii) measurements from whole nuclei (sensitized emission CM-FRET and FC-FRET) with subnuclear regions (acceptor photobleaching CM-FRET); iii) variance between CM-FRET and FC-FRET. We used transfected cells expressing either mG10, which provides a robust FRET signal at all donor/acceptor concentrations in FC-FRET or mCh+eG. Using sensitized emission CM-FRET (Figure 2A, B), cells expressing mG10 had a mean N_FRET_ of 9.9% (SD = 2.7%, n=37) while cells expressing mCh+eG had a mean N_FRET_ of 0.80% (SD = 1.09%, n=31). This is likely an underestimate of the actual FRET efficiency because sensitized emission methods are very sensitive to instrument settings and crosstalk between donor and acceptor channels. Using acceptor photobleaching CM-FRET [68] on fixed cells to eliminate diffusion into and out of the photobleached region (Figure 2C-E), cells expressing mG10 had a mean N_FRET_ of 26.8% (SD = 4.4%, n = 37) while cells expressing mCh+eG, had a mean N_FRET_ of 0.0% (SD = 2.2%, n= 31). For comparison, a typical FC-FRET experiment is shown (Figure 2G) where the mean N_FRET_ was 23.5% (SD=3.8%, n=6987), while for mCh+eG cells N_FRET_ was 0.4% (SD = 1.4%, n = 5267).

**Figure 2.**
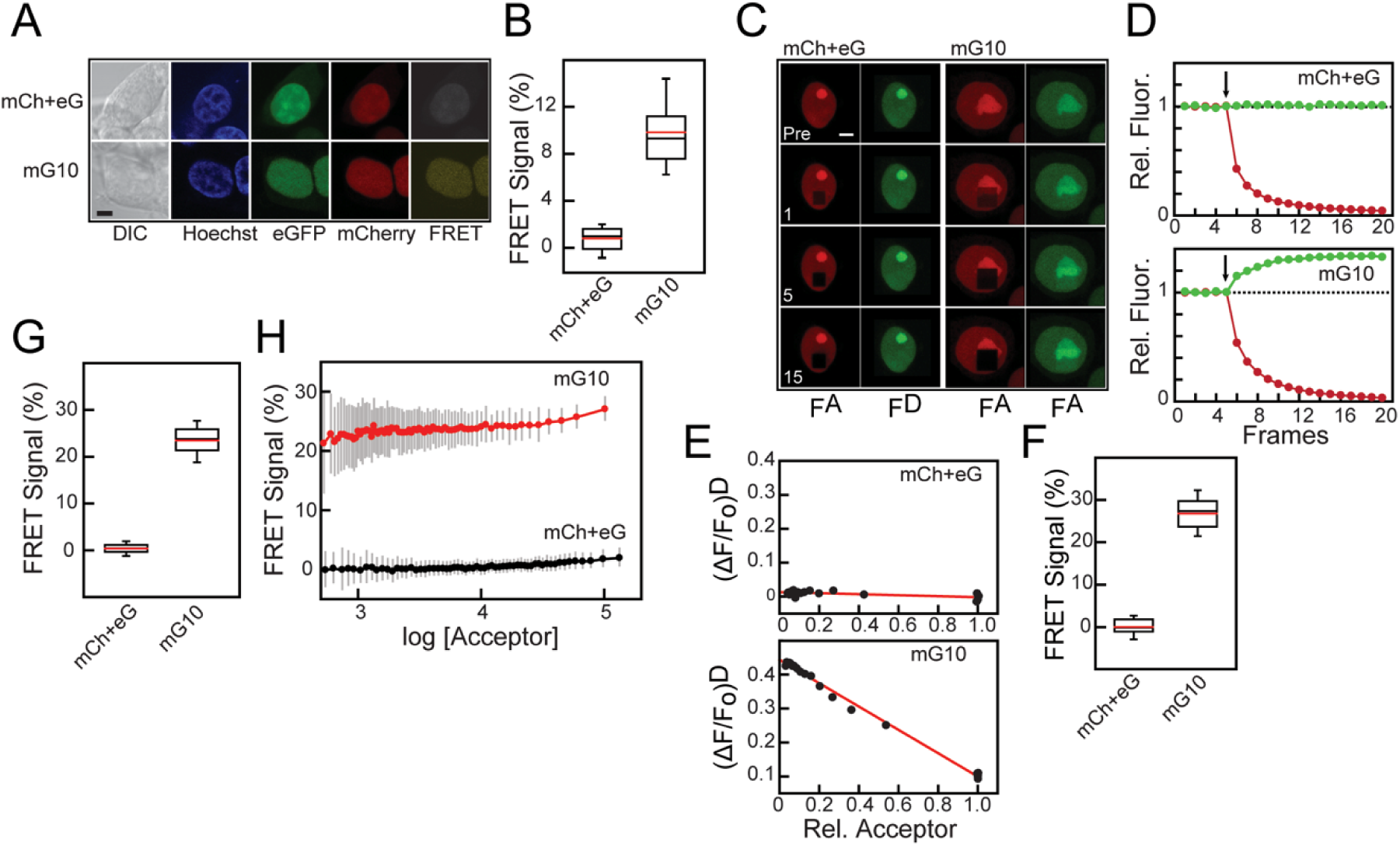
Comparison of confocal microscopy and flow cytometry FRET. A) Confocal microscopy images show HEK293 cells expressing both mCh and eG (mCh+eG, *top panels*) or an mCherry-EGFP fusion protein with a 10 amino acid linker (mG10, *bottom panels*). DIC, Hoechst (*blue*), EGFP (*green*), mCherry (*red*), FRET (*yellow*). Scale bar is 5 µm. B) Box plot showing the mean (*red line*) and median (*black line*) for sensitized emission N_FRET_ for mG10 (n=37) and mCh+eG (n=31). C) Sequential confocal microscopy images of cells expressing mCh+eG (*left panels*) and mG10 (*right panels*) before and after photobleaching. Regions were bleached after each frame with a 543 nm laser and imaged in both mCherry and EGFP channels before bleaching (*Pre*) and after 1, 5 and 15 laser pulses as. D) Fluorescence intensity changes during acceptor photobleaching in cells from (C) expressing mCh+eG (*top panel*) or mG10 (*bottom panel*). mCherry intensity (*red*) and EGFP intensity (*green*) are shown. Arrow indicates start of photobleaching. E) Plots summarizing fluorescence changes after sequential bleaching with 543 nm laser. The ordinate is the remaining acceptor fluorescence while the abscissa the fraction of donor fluorescence remaining. Measurements of relative fluorescence intensity after each bleaching laser pulse are shown (*black circles*) and the lines are a linear fit. F) Box plots showing the mean (*red lines*) and median (*black lines*) N_FRET_ from accepter photobleaching experiments for mG10 (n=37) and mCh+eG (n=31). G) FRET signals obtained in a flow cytometry experiment for individual cells expressing mG10 (*red*) or mCh+eG (*black*). Moving averages of N_FRET_ were calculated for bins of 100-cells and are plotted versus the acceptor intensity in that bin. Error bars (*grey*) are standard deviations. H) Box plots showing the mean (*red lines*) and median (*black lines*) N_FRET_ from flow cytometry experiments for cells expressing mCh+eG (n=5267) or mG10 (n=6987).

There is a wide range of expression levels in transiently transfected cells (Figures S3 and S5), but the large number of analysed cells allowed us to examine how donor or acceptor concentrations influenced FRET efficiency. The N_FRET_ for individual cells expressing mG10 or mCh+eG were plotted as a function of acceptor concentration (Figure S5). The differences in N_FRET_ were relatively constant except at the highest acceptor concentrations, where N_FRET_ from the mG10 fusion protein increased more rapidly with concentration than mCh+eG. However, the population estimates for FC-FRET represent primarily the intrinsic FRET efficiency. Moreover, the estimated N_FRET_ shows good agreement between FC-FRET and acceptor photobleaching CM-FRET efficiencies and qualitative agreement with sensitized emission CM-FRET. One possible reason for the difference in N_FRET_ measurements between FC-FRET and acceptor photobleaching CM-FRET is a potentially biased selection in the latter method of cells that have a high expression level. In addition, acceptor photobleaching CM-FRET uses a restricted subcellular region selected for measurement, with a potentially higher level of fluorescence, while FC-FRET uses the fluorescence signal from entire cell. Nevertheless, these results show that FC-FRET is a quantitative method for determining FRET efficiency in a large number of cells rapidly with comparable sensitivity to microscopic methods.

### Measurement of FRET between CRX donor and acceptor

We used flow cytometry to examine potential FRET between individual human CRX molecules with terminal fusions with mCherry (m) or EGFP (e) (Figure 3A) expressed in HEK293T cells. To ensure that nuclear localization was not disrupted by N-terminal fusions, a nuclear localization signal was added to these constructs at the beginning of either mCherry or EGFP. Thus, subcellular localization assessment of the native proteins is confounded by this modification. However, the purpose of the present study was to examine interaction of the CRX and NRL in the nucleus. CRX (see Figure 3 legend for nomenclature) distributed exclusively in the nucleus with a nonhomogeneous distribution (Figure 3B shows representative images). All CRX fusions showed similar subnuclear distributions (*data not shown*). N_FRET_ was highest (3.3%, p<0.001, one way ANOVA, Holm-Sidak, pairwise comparison with all other groups) when both donor and acceptor were fused to the N-terminus near the homeodomain (Figure 3C). N_FRET_ was lower when the fusion proteins were on the C-terminus following the activation domain and not significantly different compared to mCh+eG (1.6%, p=0.452, one way ANOVA, Holm-Sidak). N_FRET_ was at background levels comparable with mCh+eG when the fusion proteins were on different termini (CRXm+eCRX, p=0.828 and mCRX+CRXe, p=0.917, one way ANOVA, Holm-Sidak). Cells transfected with CRX fusion proteins together with a soluble fluorescent protein also exhibited background FRET signals (Figure 3C).

**Figure 3.**
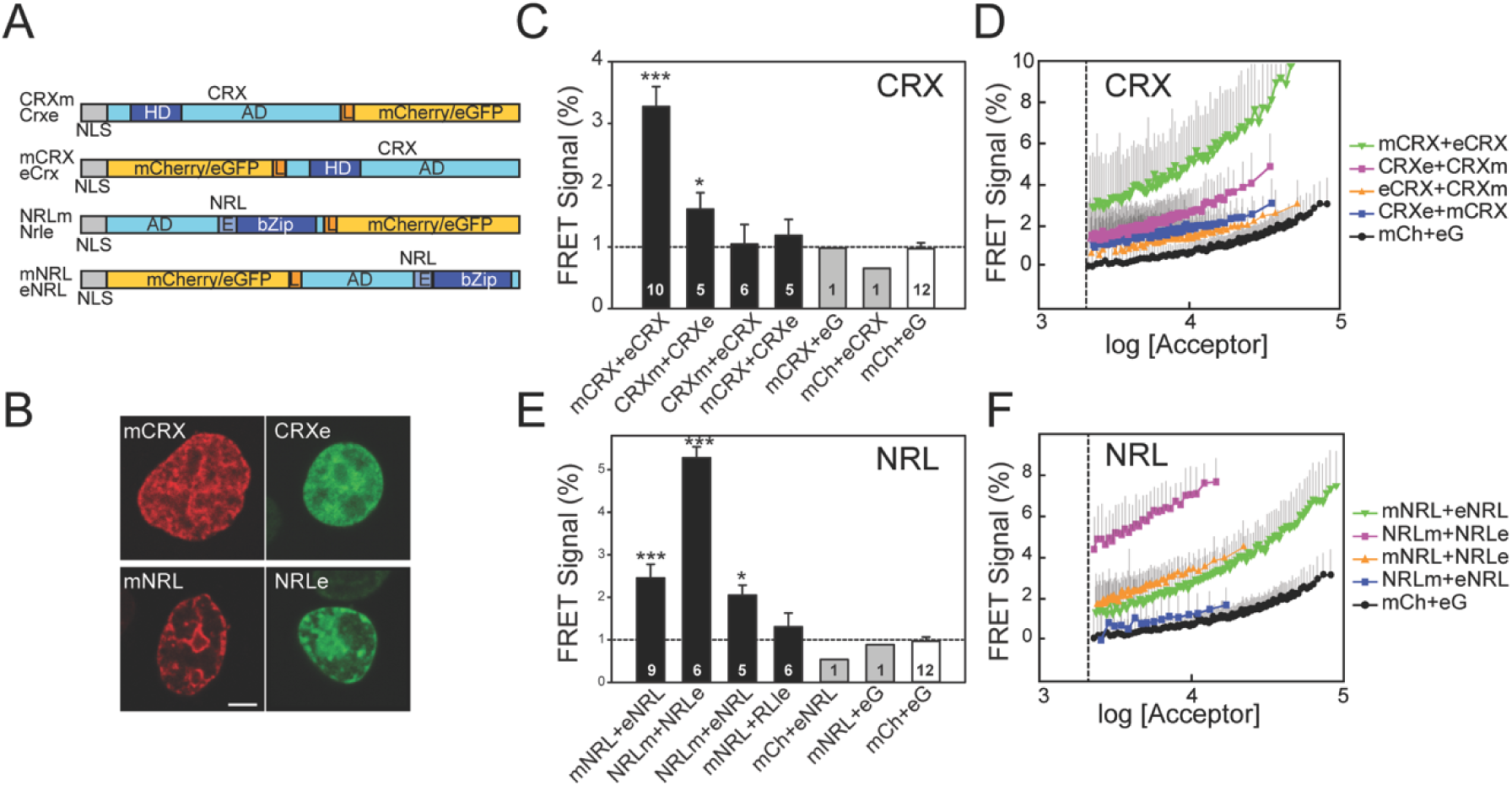
FRET between fluorescently tagged CRX and NRL homo-pairs. A) Fluorescent CRX and NRL constructs used for FRET analysis. Diagram illustrates the fusions between fluorophores (mCherry and EGFP) and at the N and C-termini of CRX or NRL. All constructs have an N-terminal nuclear location signal (NLS). Domains of CRX and NRL are indicated: activation domains (AD), homeodomain (HD), basic leucine zipper domain (bZip), and linkers (L). The constructs are labeled with the fluorophore (m: mCherry or e: EGFP) at the beginning or end of the label depending upon the terminus to which it is fused. For example, mNRL is NRL with mCherry fused to the N-terminus. B) Confocal microscopy images of transiently transfected HEK293T cells expressing a fluorescently tagged transcription factor as indicated. Cells had almost exclusively nuclear fluorescence, with a patchy intranuclear pattern. Scale bar is 5 µm. C, E) Comparison of FRET signals determined by flow cytometry using HEK293T cells cotransfected with combinations of donor-acceptor CRX (C) or NRL (E) fusion constructs as indicated in the panel. The bars represent mean values with standard errors of the FRET signals from the indicated number of flow cytometry experiments. For each experiment, more than 1000 cells were analyzed. Statistical significance was performed using ANOVA (compared with mCh+eG cells): p<0.05 (*) and p<0.001 (***). The samples with only a single flow cytometry experiment were not used in the ANOVA. The dotted line indicated the mean FRET signal for mCh+eG. D,F) FRET signals in individual cells expressing combinations of donor and acceptor fused to CRX (D) or NRL (F) are plotted as a function of acceptor fluorescence intensity. Symbols are the mean FRET signal in each bin (100 cells) and lines are moving averages. Error bars (*grey*) are the SD, only the positive SD is plotted for clarity. Cells with fluorescence values below threshold fluorescence intensity (*dashed line*) were not included in the analysis.

We examined the FRET signal from the various CRX fusion proteins as a function of expression level. N_FRET_ was significantly higher at all fluorescence levels when both donor and acceptor were fused to the N-terminus near the homeodomain (mCRX+eCRX) in comparison to all other combinations (Figure 3D). The slope for the mCRX+eCRX pair increased much faster as expression levels increased compared to mCh+eG or pairs with fluorophores on opposite termini (eCRX+CRXm and CRXe+mCRX), which were similar to mCh+eG at all concentrations. N_FRET_ for constructs with fluorophores fused to the C-terminus near the activation domain (CRXe+CRXm) were intermediate between mCRX+eCRX, showing elevated N_FRET_ at higher expression levels than expected from stochastic FRET alone [59]. These data indicate that a fraction of CRX molecules expressed in HEK293T cells are close enough for FRET, with fusions having both donor and acceptor near the homeodomain giving larger FRET signals than with fusions both near the activation domains. These results suggest that the FRET-detectable fraction of CRX molecules is arranged in a head to head fashion.

### Measurement of FRET between NRL donor and acceptor

We used flow cytometry to examine potential FRET between individual human NRL molecules with terminal fusions with mCherry or EGFP (Figure 3A) expressed in HEK293T cells. NRL fusions were distributed exclusively in the nucleus and had nonhomogeneous distributions (Figure 3B shows a representative image). All NRL fusions showed similar subnuclear distributions (*data not shown*). N_FRET_ was highest (5.3%) and significantly different from all other groups (p<0.001, one way ANOVA, Holm-Sidak, pairwise comparison) when both donor and acceptor were fused to the C-terminal bZIP domain (Figure 3C). N_FRET_ was lower (2.5%) when the fluorescent proteins were both fused to the N-terminus but was significantly different than mCh+eG (p<0.001, one way ANOVA Holm-Sidak). N_FRET_ was at background levels for NRLm+eNRL (p=0.349, one way ANOVA, Holm-Sidak, compare with mCh+eG) but slightly higher for mNRL+NRLe (2.1%, p=0.013, one way ANOVA, Holm-Sidak). This may reflect differences in relative angles between the fluorophores in the two complementary pairs. Control experiments in which cells were transfected with an NRL-fusion protein and a soluble fluorescence protein (mCherry or EGFP) exhibited N_FRET_ comparable to mCh+eG (Figure 3C).

We examined the FRET signal from the various NRL fusion proteins as a function of expression level. Fusions to the C-terminus, near the bZip DNA binding domain, did not give expression levels as high as fusions to the N-terminus or for CRX fusion proteins (compare Figures 3D and 3F). However, N_FRET_ was much higher at all fluorescence levels when both donor and acceptor were fused to the C-terminus (NRLm+NRLe) in comparison to all other combinations (Figure 3F). The slope for the NRLm+NRLe pair increased faster as expression levels increased compared to mCh+eG. N_FRET_ from constructs with fluorophores fused to the N-terminus near the activation domain (mNRL+eNRL) also increased with expression level much faster than mCh+eG (Figure 3F). When fluorophores were fused to opposite termini (mNRL+NRLe, and NRLm+eNRL), N_FRET_ behaviour diverged for reasons that are not clear. The mNRL+NRLe pair had similar N_FRET_ compared to mNRL+eNRL pair in the overlapping expression range, which was higher than mCh+eG. The other combination, NRLm+eNRL, was similar to mCh+eG. The elevated N_FRET_ for three of the four NRL combinations, particularly NRLm+NRLe, were higher than expected from stochastic FRET alone [59]. These data indicate that a fraction of NRL molecules expressed in HEK293T cells are close enough for FRET, with fusions having both donor and acceptor near the DNA binding domain giving larger FRET signals than with fusions both near the activation domains. These results suggest that the FRET-detectable fraction of NRL molecules is arranged in a tail to tail fashion. The FRET efficiency for the C-terminal FRET pair is similar to that observed (∼ 4%) for a heterodimer of Fos and Jun, both of which are bZIP proteins [51].

### Measurement of FRET between CRX and NRL donor-acceptor pairs

We used flow cytometry to examine potential FRET between individual CRX and NRL molecules with terminal fusions with mCherry or EGFP (Figure 3A) expressed in HEK293T cells. The nuclear distribution pattern for both CRX and NRL when expressed in the same cell were similar but not identical (Figure 3B). We observed FRET between CRX and NRL with a number of donor-acceptor fusion pairs (Figure 4A). Fusions at the N-termini of NRL and CRX had the highest N_FRET_ compared to mCh+eG (p=0.001, one way ANOVA, Holm-Sidak). Fusions with fluorophores both located at the C-termini of CRX and NRL had N_FRET_ higher than mCh+eG but it was not statistically significant (p=0.45, one way ANOVA, Holm-Sidak). Constructs with donor and acceptor on opposite termini were not different from mCh+eG (Figure 4A).

**Figure 4.**
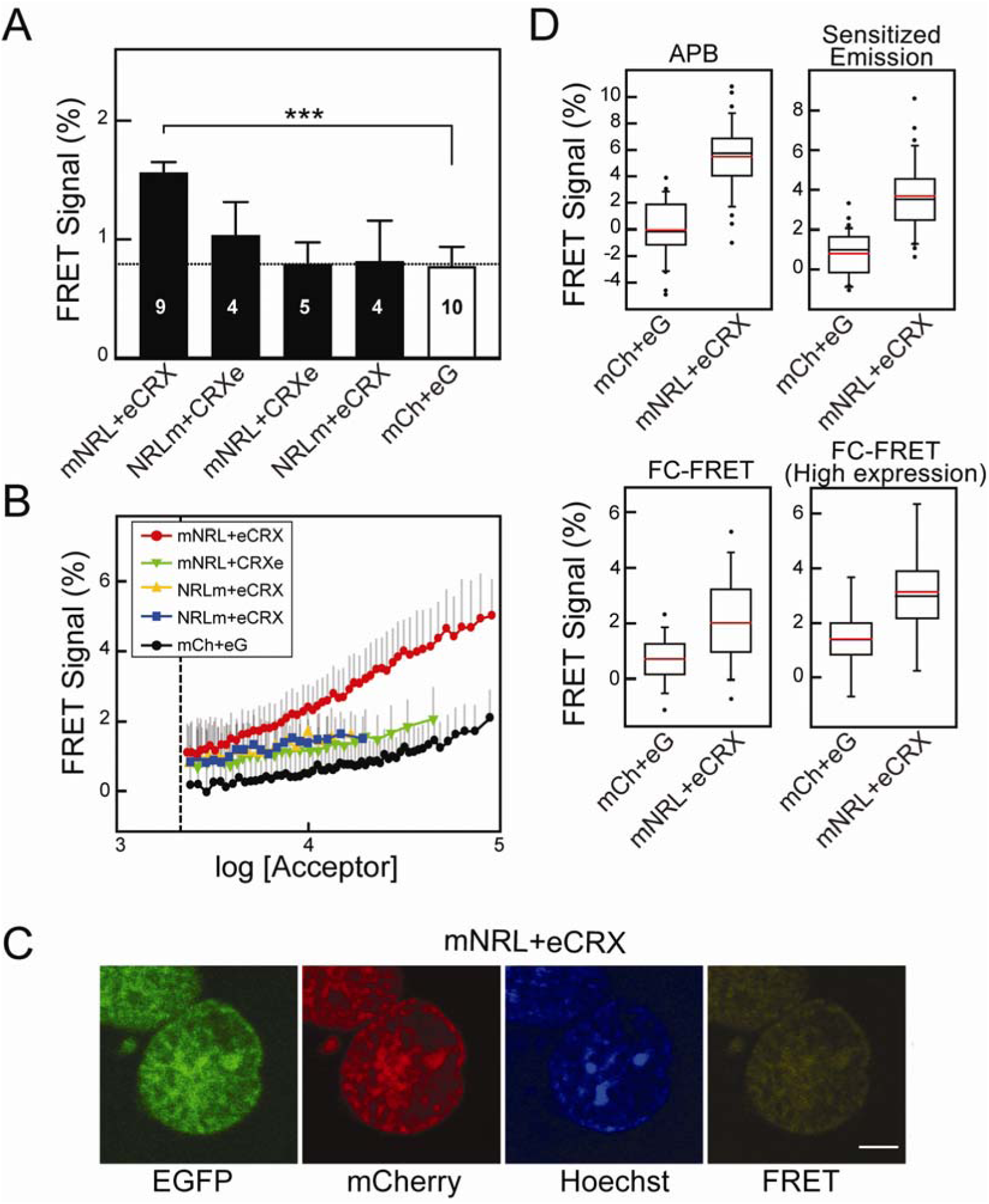
FRET between fluorescently tagged CRX and NRL hetero-pairs. A) Comparisons of FRET signals determined by flow cytometry using HEK293T cells cotransfected with combinations of donor-acceptor CRX and NRL fusion constructs as indicated. The bars represent mean N_FRET_ with standard errors from the indicated number of flow cytometry experiments indicated. For each experiment, more than 1000 cells were analyzed. The dotted line indicates the mean FRET signal for mCh+eG. Statistical significance was determined by ANOVA and pairwise comparisons with mCh+eG cells: p<0.001 (***). B) FRET signals in individual cells expressing combinations of donor and acceptor fused to CRX and NRL as indicated are plotted as a function of acceptor fluorescence intensity. Symbols are the mean FRET signal in each bin (100 cells) and lines are moving averages. Error bars (*grey*) are the SD, only the positive SD is plotted for clarity. Cells with fluorescence values below threshold fluorescence intensity (*dashed line*) were not included in the analysis. C) Confocal microscopy images of the nuclei of transiently transfected HEK293T cells co-expressing fluorescently tagged NRL (mNrl) and CRX (eCrx). Fluorescence was recorded in EGFP, mCherry, and FRET channels. Hoechst was used label DNA. Scale bar is 5 µm. D) Box plot comparisons of FRET signals for CRX-NRL donor and acceptors measured by microscopy-based FRET (*acceptor photobleaching (APB)* and *Sensitized Emission*) and one flow cytometry FRET experiment. The numbers of cells analyzed (mCh+eG/mNRL+eCRX): APB, n = 31/34; Sensitized Emission, n = 31/34; FC-FRET, n = 6996/8237; FC-FRET (High-expression, F^A^ >10^4^), n = 1031/953. Mean (*red lines*) and median (*black lines*) N_FRET_ are indicated. Statistical significance for each comparison was significant by t-test with P<0.001 (***).

We examined FRET from the various combinations of fusion proteins as a function of expression level (Figure 4B). N_FRET_ from constructs with both donor and acceptor fused to the N-terminus (mNRL+eCRX) was statistically higher than mCh+eG at all fluorescence levels and than the other combinations when F^A^ >10^4^. The slope for the mNRL+eCRX pair increased faster as expression levels increased compared to mCh+eG, suggesting a possible concentration dependence on the proximity of mNRL and eCRX. The FRET signals for the other fusion constructs increased with expression level similar to mCh+eG. What is clear from the dependence on expression level is that the FRET signal from mNRL+eCRX is much higher than expected from stochastic FRET alone [59]. These data indicate that a fraction of NRL and CRX molecules when expressed in HEK293T cells are close enough to FRET, with fusions having donor near the DNA binding domain of CRX and acceptor near the activation domain of NRL giving large FRET signals.

We compared N_FRET_ between mCRX and eNRL using FC-FRET to those measured with CM-FRET (Figure 4C). Both acceptor photobleaching and sensitized emission approaches had significant N_FRET_ for N-terminal fused mNRL+eCRX when compared to mCh+eG (t-test, p<0.001, respectively). These data agree with the results from the FC-FRET experiments. The CM-FRET methods gave a higher estimated efficiency because the measurements were performed on nuclear regions with high fluorescence levels where both donor and acceptor overlap, while the FC-FRET measurements are derived from the entire nucleus, which may underestimate locally confined interactions. Taken together, these results confirm that a fraction of CRX and NRL molecules in HEK293T nuclei are close enough for FRET.

## Discussion

We report improvements in the use of flow cytometry to obtain FRET efficiencies. The major advantages are the large number of cells and the wide-range of donor-acceptor expression levels available for N_FRET_ determinations. We used this method to characterize CRX and NRL dimer interactions in live cells. In order to investigate homodimer interactions, we measured FRET signals between donor and fusion pairs from the same transcription factor co-expressed in individual cells. In addition, we detected formation of CRX-NRL complexes in live nuclei for the first time.

NRL with donor and acceptors fused to the C-terminus (adjacent to the bZip domain) gave the largest average FRET signals. Fusions at the N-terminus (adjacent to the activation domain) also gave robust FRET signals, but they were lower than the C-terminal fusions. These results indicate that a fraction of NRL donor and acceptor fusion proteins, when expressed in the same cell, are close enough to generate a large FRET signal. When donors and acceptors were near the DNA binding domain (bZip) the FRET signals were larger than when placed adjacent to the activation domain. These results are consistent with a parallel orientation (tail to tail) of the NRL dimer (Figure 5A). If a simplified scenario is assumed, it is possible to make an estimate of the proximity of donor-acceptor NRL dimers. First, all of the expressed NRL is assumed to be involved in FRET interactions and thus the use of the total fluorescence intensity in the N_FRET_ calculation would be justified. Second, the average donor-acceptor orientations are assumed to be similar in the mG fusion series. With these two assumptions, estimated proximity can be derived from the calibration of N_FRET_ from the mG fusion series (Figure 2E). With this ruler, the FC-FRET data implies a donor-acceptor distance at the C-terminal (bZip) region between NRL monomers ∼6-7 nm apart and ∼10 nm for the N-terminal region. Future experiments are needed to investigate the proportion of donor and acceptor molecules that that are interacting. In addition, we note that with these assumptions, the estimated distance is a conservative one, since the actual FRET efficiency is probably higher than determined in the FC-FRET measurements. NRL is part of the large MAF family and forms homo-and heterodimers with other bZip transcription factors [35]. Therefore, we expect that NRL will not be monomeric in HEK293T cells, rather it could be in complexes with itself or other other MAF or bZip proteins proteins expressed in HEK293T cells [69]. Nonetheless, the distance estimate from the N_FRET_, measurements seem reasonable given the following distance estimates: 1) the structure of the closely related MAFA (DNA binding domains for the MAFA dimer bound to DNA are ∼1 nm apart, PDB ID: 4EOT [70]), 2) the diameter of donor-acceptor molecules (∼2.5 nm, [71]) and 3) the length of the (Gly)_5_ linkers between the fluorophores and NRL (∼2 nm). The estimates further support the identification of the N_FRET_ measure with FC-FRET as a true proximity indicator.

**Figure 5.**
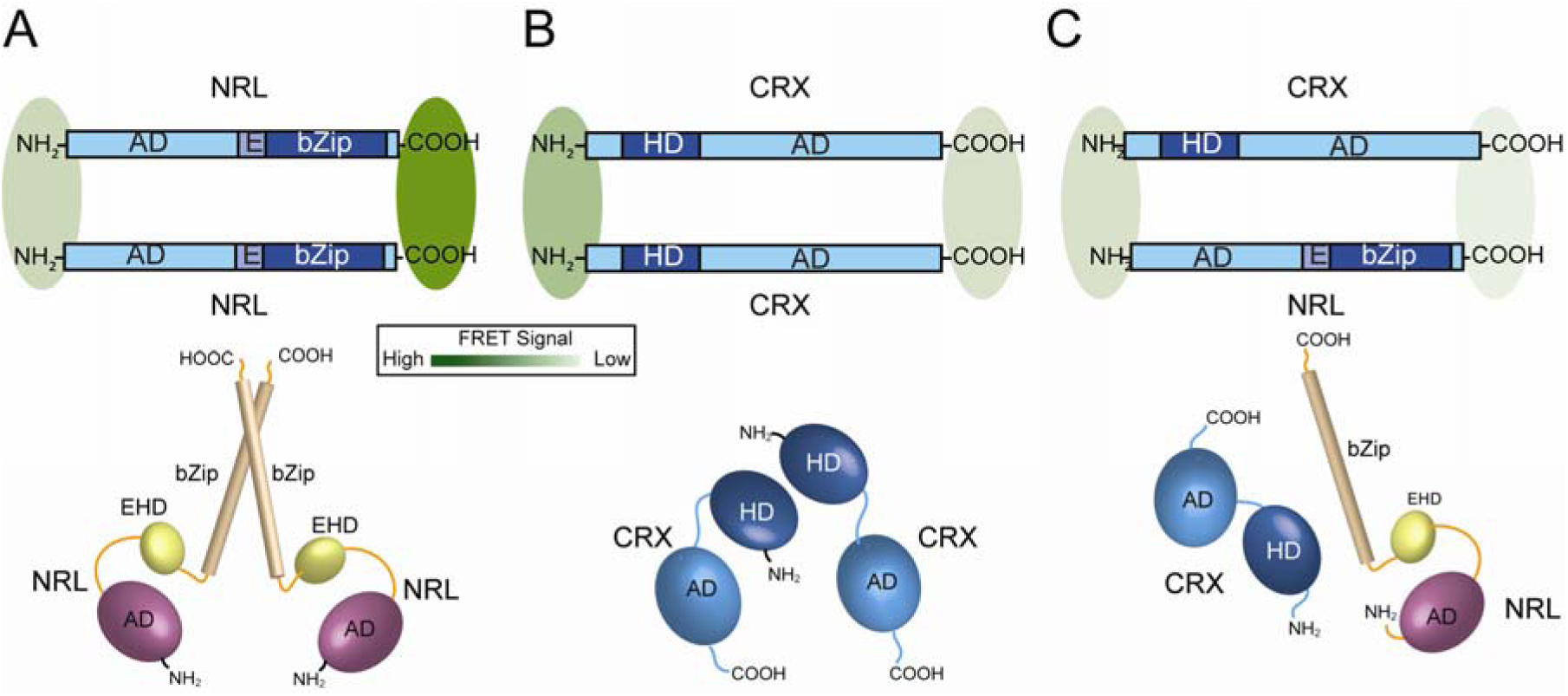
Models of NRL and CRX interactions based on FRET signals. *Upper Diagrams*. Schematic diagrams illustrating possible arrangements of NRL (A) and CRX (B) homodimers and NRL-CRX heterodimer (C). The intensity of the FRET signals obtained for the various combinations (Figures 4 and 5) are illustrated as green shaded ovals, with the intensity of the color representing the relative FRET signals for the various constructs. *Lower Diagrams*: Speculative three-dimensional arrangement of the various domains in NRL and CRX complexes.

Crx with donor and acceptors fused to the N-terminus (adjacent to the HD) gave a large average FRET signal but less than obtained with Nrlm/Nrle. Fusions at the C-terminus (adjacent to the activation domain) also gave FRET signals that were statistically above control levels, but they were lower than the N-terminal fusions or those obtained with Nrl. These results indicate that a fraction of Crx donor and acceptor fusion proteins, when expressed in the same cell, are close enough to generate a large FRET signal. These results are consistent with a parallel orientation (head to head) of the CRX dimer (Figure 5B). Assuming again the simplified scenario for estimating proximity, the FC-FRET data implies a donor-acceptor distance at the N-terminal (HD) region between CRX monomers of ∼10 nm. Again, future work is needed to further investigate proportion of molecules involved in dimerization. The formation of of Crx dimers is consistent with observations on other members of the *paired* homeodomain family [72-75]. Together, the FC-FRET results support the conclusion that both Crx and Nrl can assemble into homodimers (or oligomers) in live cells. Moreover, placing donors and acceptors near the DNA binding domains gave the largest FRET signals. These are the first data to our knowledge that reveal oligomeric organization of either CRX or NRL.

In the case of CRX and NRL, the N_FRET_ was smaller than for the homodimers, but significantly greater than mCh+eG controls. Moreover, N_FRET_ steadily increased as expression levels increased (Figure 4B), at a rate much faster than observed for stochastic FRET. However, the robust FRET signals from CRX and NRL donor-acceptor pairs, distinctly higher than control donor-acceptor pairs, demonstrate formation of CRX-NRL complexes in live nuclei for the first time. This data is consistent with biochemical [36] and functional data [7, 36, 37] that show their interaction suggests possible mechanisms for their cooperative activity. Because of the small FRET signal, we are not able to estimate proximity between the donor and acceptors fused to CRX-NRL dimers. It is important to emphasize that we do not know what fraction of donor or acceptors expressed in the nuclei of HEK293T are available to interact or potentially participate in FRET. We are not able to determine that FRET fraction using CM-FRET approaches – additional studies require techniques such as single molecule FRET [76, 77].

There are several advantages in the use of this improved FC-FRET technique over previous applications [45-53]. Here, we showed that FC-FRET is sensitive enough to detect subtle differences in donor-acceptor linker lengths, and thus distance. In addition, we have a standardized data acquisition and analysis workflow. In this implementation, there is a delay between exposure of cells to donor (488 nm) and acceptor (561 nm) excitation lasers, thus avoiding crosstalk between fluorescent proteins in the detection channels during their fluorescence lifetimes. FC-FRET employs one of the most widely used sensitized emission methods, called N_FRET_ [78], to calculate FRET efficiency. N_FRET_ minimizes the dependence of FRET efficiency on the donor and acceptor fluorescence intensities. However, N_FRET_ deviates dramatically from expected behaviour when the stoichiometry of donor or acceptor are not matched well [44]. FC-FRET can address this issue by the selection of cells with a desired acceptor/donor ratio where N_FRET_ is stable [58], here between 0.1 - 10 range. FC-FRET allows a direct examination across the entire range of fluorophore expression, permitting a qualitative separation of the FRET signal into intrinsic and stochastic components by comparison with co-expressed, unlinked donor and acceptor. This is particularly important when FRET efficiency between interacting proteins is low, because of distance between donor/acceptor or orientation of the donor/acceptor; or because the fraction of interacting molecules is low (for example, due to competing cellular binding partners or influence of the K_D_ on extent of interaction), so the FRET signal, which is normalized to the total donor/acceptor concentrations, is artificially low.

The FC-FRET results quantitatively agree with measurements made using two common microscopy-based methods and are comparable between different transfections and flow cytometry runs. FC-FRET analysis uses individual cell FRET efficiencies for statistical analysis and includes information on donor and acceptor fluorescence intensities. Thus, FC-FRET efficiencies are obtained across a wide range of expression levels. This allows the contribution of stochastic FRET to be accounted for directly. FRET efficiency is determined from total cellular fluorescence, which potentially reduces bias in the (subjective) collection of cells with intense fluorescence. In summary, we have shown that FC-FRET is able to quantitatively study protein-protein interactions in live cells in a high-throughput manner.

Although FC-FRET has advantages, it shares several limitations with whole-cell microscopy and solution methods. FC-FRET does not provide detailed information on subcellular distribution, only measures average fluorescence intensity for each cell. If a donor and acceptor are concentrated in a particular location, the stochastic FRET efficiency could be higher than expected and thus be mistaken for protein-protein interactions. Therefore, it is imperative to design control fluorescent proteins that appropriately colocalize and produce the same expression levels as the candidate proteins for comparison. FC-FRET sensitivity is limited by the fraction of donor and acceptor that can interact with each other. For example, if the donor and acceptor only interact in certain compartments, but are also found in other compartments, then the apparent FRET efficiency will be reduced. A strategy that combines microscopy and flow cytometry would overcome this limitation.

In conclusion, we have developed FC-FRET, an improved flow cytometry-based FRET method that validated using confocal microscopy FRET methods. The most significant advantage is the ability to analyze FRET signals from cells with a wide range of expression levels. The permits a separation of the FRET signal into intrinsic and stochastic components. Moreover, we have shown for the first time an interaction between CRX and NRL in live cells. Using this approach, we have proposed orientations between NRL and CRX homodimers/heterodimers.

## Methods

### Expression constructs

Fusion constructs and large deletions were generated by overhang extension PCR [79] using primers from IDT (IDT, Coralville, IA) and cloned Pfu DNA polymerase (Stratagene, La Jolla, CA). Point mutations and small deletions or insertions were generated using QuickChange [80] with Turbo Pfu DNA polymerase (Stratagene, La Jolla, CA). A nuclear localization signal (NLS), MAPKKKRKVNRSKA, was added at the N-termini of EGFP and mCherry (Clontech, Mountain View, CA). For intramolecular FRET experiments, EGFP and mCherry fusion proteins (mG) were designed with various linkers (Supplemental Table 1). The α-helical linkers were based on a repeated (n=2-7) α-helix-forming peptide, EAAAK (27) flanked by two proline residues to terminate the α-helical region. For expression of NRL and CRX fusion proteins, coding regions were cloned downstream of the CMV promoter in derivatives of the pEGFP-N1 plasmid (Clontech) with an NLS and linker sequences to EGFP or mCherry (Supplemental Table 2). All constructs were confirmed by DNA sequencing (Genewiz, www.genewiz.com).

### Mammalian cell culture and transfection

HEK293T cells (ATCC, Manassas, VA) were cultured in DMEM supplemented with 10% FBS and 1 mM L-glutamine. Cells were seeded at 75,000 cells/ml one day before transfection. Cells were transfected with a total of 1 µg of DNA using Fugene 6 (Roche, Branchburg, NJ) according to the manufacturer’s instructions

### Confocal Microscopy FRET

In sensitized emission FRET and live cell imaging experiments, HEK293T cells were seeded on a collagen coated No. 1 coverslip placed in the bottom of a 3.5 cm dish (MatTek, Ashland, MA) before transfection. One day after transfection, cells were placed in phenol red-free DMEM (Gibco, Carlsbad, CA) containing 0.1 µg/ml Hoechst 33342 (Sigma-Aldrich, St. Louis, MO), 10% FBS and 1 mM L-glutamine and incubated for one hour. Cells were then placed in the environmental chamber (PeCon GmbH, Germany) of the confocal microscope in 5% CO_2_ at 37°C and equilibrated for 15 min. Confocal images were collected using a LSM510 META microscope (Carl Zeiss, Germany) equipped with a Plan-Apochromat 63× oil immersion objective (NA 1.4) and an Argon laser (488 nm) and a HeNe laser (543 nm). The pinhole was adjusted to obtain 1 Airy unit for the 488 nm laser. To reduce contamination signals between the two fluorescence channels, 500-535 nm band pass and 560LP long pass filters were used to filter fluorescence excited by Ar and HeNe lasers, respectively. The FRET signal was detected using the Argon laser 488 nm line and a 560LP long pass filter. Hoechst 33342 staining was detected using a two-photon Chameleon laser exciting at 800 nm (power 4-8%) and a 435/485 nm band pass filter. For dual color acquisition, 12-bit images were sequentially acquired in a line-scan mode (average of two scans). The images were filtered by one-time Gaussian blur (0.5 sigma) in ImageJ (NIH) to reduce noise. The fluorescence intensity for sensitized FRET analysis was quantified from the filtered images (N_FRET_, details described in FC-FRET section). For presentation in the figures, filtered image brightness and contrast were adjusted using ImageJ for the entire image. In APB FRET, HEK293T cells were seeded in 8 well chamber slides (Nunc Lab-Tek), transfected as described above and then fixed with 2% paraformaldehyde for 15 min. Slides were mounted in glycerol prior to image acquisition as described above. Acceptor was sequentially photobleached using the HeNe laser at 100% power. Images were analyzed using Image J software (NIH) and Sigma Plot 11.0 (Systat Software, Inc., Chicago, IL).

### Flow cytometry FRET

Cells were transfected as described above, treated with 0.1% trypsin for 5 min and then washed in phenol red-free DMEM containing 10% FBS. Cells were centrifuged at 250 g for 5 min and suspended with phosphate buffered saline at ∼10^6^ cells/ml. FC-FRET measurements were performed using a LSRII flow cytometer (BD Bioscience) equipped with 405 nm, 488 nm, 561 nm and 633 nm lasers. A 19 µs delay was set between 488 nm and 561 nm laser interrogation times. To measure EGFP and FRET fluorescence intensities, cells were excited with the 488 nm laser line with fluorescence collected in the EGFP channel through a 530/30 band pass filter, while the FRET signal was collected through a 610/20 band pass filter. To measure mCherry fluorescence, cells were excited with the 561 nm laser line and fluorescence was collected through a 610/20 band pass filter. Channel settings were optimized and calibrated as follows. First, the voltage of each photomultiplier was adjusted to balance the fluorescence intensity for EGFP and mCherry. Second, the fluorescence intensity for each channel was calibrated with beads having known amounts of fluorophore attached (Spherotech, Inc.). Finally, the FSC (forward scattering) and SSC (side scattering) were calibrated with beads of known size (Spherotech, Inc). The concentration of fluorescence molecules was estimated by the fluorescence intensity and estimated size of each cells. We calibrated the fluorescence intensity and size measurement with Spherotech beads as described above. We converted intensity to equivalent brightness of fluorescence dyes (such as EGFP). We use this number to estimate the number of fluorescence molecules in a cell. We used the size standard beads to estimate the size of cells with FSC and SSC reading. We assume a HEK293 cell can approximate to a spherical ball in a solution. Based on this assumption, we estimate the volume of each cell and then the concentration of fluorescence molecules based on the number of fluorescence molecules in a cell.

For each experiment, four control groups were analysed. Mock transfected cells were used to set background fluorescence levels for donor, FRET and acceptor channels. Cells expressing only EGFP were used to measure the bleed-through of donor emission (EGFP) into the FRET channel (610/20), calculated as the ratio of donor emission detected in the FRET (acceptor) channel to donor channel, D_C_. Similarly, cells expressing only mCherry were used to measure the excitation of acceptor (mCherry) by donor excitation light (488 nm), calculated as the ratio of acceptor emission with donor excitation to acceptor fluorescence, A_C_. The variation in D_C_ and A_C_ between cells decreased as both acceptor and donor fluorescence intensity increased, respectively, and we used a value of 30% variation in D_C_ and A_C_ to set the lower limit of EGFP and mCherry intensities for including cells in the analysis. We used a sensitized emission calculation, also called the three-cube method [55], to determine the normalized FRET efficiency in FC-FRET:

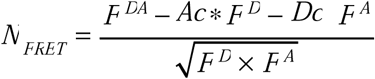

where F^DA^, F^D^and F^A^ are the fluorescence intensities in the FRET, donor and acceptor channels, respectively. Flow cytometry data files were imported to custom software for data processing in the Matlab (Mathworks, Inc) environment (executable program available upon request). Different group mean or median values were compared with Student’s t-test (if normality test failed, a Mann-Whitney Rank Sum test was used) or ANOVA analysis (Holm-Sidak method) in Sigma Plot 11.0 (Systat Software, Inc., Chicago, IL) using p<0.05(*), p<0.01(**) and p<0.001(***).

### Simulation of stochastic (collisional) FRET

To simulate the effect of concentration on unlinked donor-acceptor fluorophores, we used the approach as described by Lakowicz for freely diffusing donor-acceptor pairs [59-61]. To calculate the FRET efficiency for collisional events we used 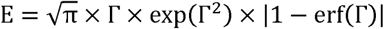, where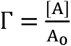 is the ratio of effective acceptor concentration to the critical concentration A_0_, which represents the acceptor concentration that results in 76% energy transfer. In the case of mCherry and EGFP, we calculated the stochastic FRET to be greater than ∼1% when the concentrations of mCherry and EGFP are higher than ∼20 µM (Figure S8A). The result of the acceptor concentration dependent FRET measurement with mCherry and EGFP is close to this value (∼10 µM), suggesting the FRET efficiency above this level will have a stochastic FRET component. However, this analysis of stochastic FRET does not include an exclusion volume for large fluorophores. To include that variable in simulations of a three dimensional collisional system, we used a Monte Carlo approach based on a randomized static distribution of acceptors and donors (Figure S8B) to calculate the FRET efficiency by proximity using the Förster equation and summing over the closest pairs. In this simulation, we used κ^2^ = 0.476 (instead of 2/3), resulting in an energy transfer for EGFP-mCherry at the critical concentration, A_0_ = 3.6 mM, to be 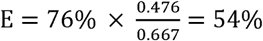. The simulations (using in MatLab (Natick, MA), R2012b) were performed over a range of concentrations (total molecules 50-600) in volumes of spheres with radii of 100-600 nm and differing distance constraints (3-6 nm) for the FRET efficiency calculation. A total of 21 random distributions were used to generate Figure S8C.

## Acknowledgments

This work was supported in part by the National Institutes of Health grants EY-11256 and EY-12975 (B.E.K.) and the Research to Prevent Blindness (Unrestricted Grant to SUNY UMU Department of Ophthalmology). We thank Dr. S. Reks critical comments on this work as it progressed.

## Author Contributions statement

X.Z and B.E.K. designed research; X.Z. performed research; X.Z and B.E.K. analyzed data; and X.Z and B.E.K. wrote the paper. Both authors reviewed the manuscript.

