## Supplemental Data for "Interaction of human CRX and NRL in live HEK293T cells measured using fluorescence resonance energy transfer (FRET)"

### Supplementary Figures and Tables

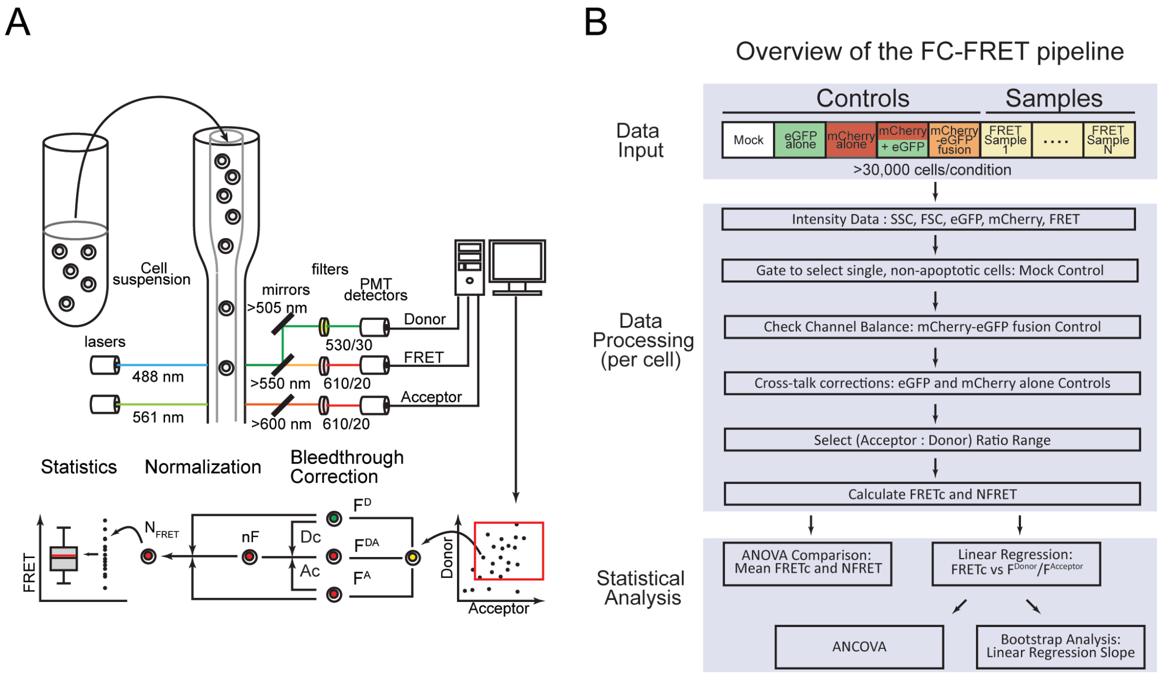


**Figure S1. Flow cytometry to detect FRET signals.**
The experiment setup (A) and data processing workflow (B) of flow cytometry-based FRET. Live HEK293T cells were transfected with nuclear localized mCherry-eGFP fusion proteins (mG) or transcription factors tagged with either mCherry or eGFP. Cells were analysed on a BD LSRII flow cytometer. Cell were transported in a sheath fluid and streamed to the laser beams for interrogation. Two lasers, 488 nm and 561 nm were used in this system with different focusing position. The 488nm laser excited eGFP and FRET signals, which were directed through 530/30 and 610/20 optical filters to the donor ($F^D$) and FRET PMT Channels ($F^{DA}$) accordingly. The 561 nm laser excited the mCherry signal that was directed through 610/20 optical filters to the acceptor PMT Channel ($F^A$). The FRET signal ($F^{DA}$) was corrected for background fluorescence, spill-over from donor (Dc) and excitation of acceptor (Ac). The corrected FRET signal was normalized with donor and acceptor signal to generate the FRET signal ($N_{FRET}$). The cell by cell FRET result then underwent further desired statistical analysis.

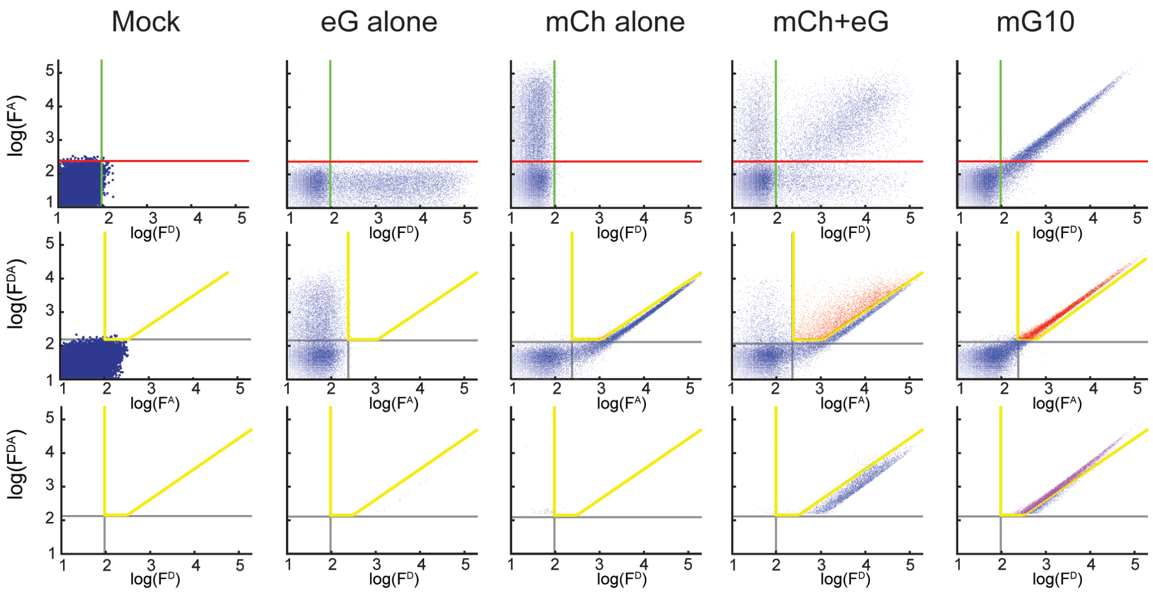


**Figure S2. Fluorescence intensity distributions in HEK293T cells transfected with fluorophores analysed by multichannel flow cytometry.**

Distributions of the five cell populations used in FC-FRET analysis: eGFP ($F^D$), mCherry ($F^A$) and FRET ($F^{DA}$) fluorescence intensities. The threshold levels for eGFP (*green*) and mCherry (*red*) fluorescence channels are shown in the top row. The FRET threshold levels after correcting for background, spill over and cross-talk are shown (yellow). Cells that cross over the FRET threshold are shown in red. Although the cell number is readily determined for the various samples, it is not possible to determine a FRET efficiency with cell counts alone. This limits the sensitivity of flow cytometry methods based on this strategy.

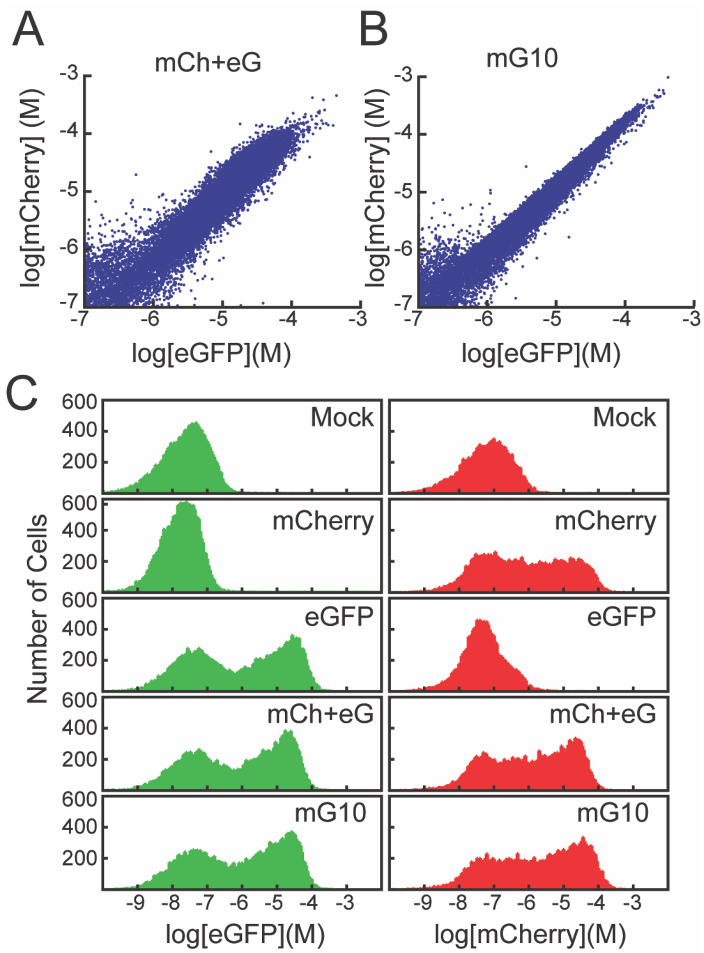


**Figure S3. Expression levels and fluorescence intensities in transfected HEK293T cells.** A, B) Scatter plots of mCherry (red) and eGFP (green) concentration in HEK293T cells transfected with mCh+eG (A) and mG10 (B). Concentrations were determined from the flow data as described in Methods. C) Histograms of mCherry and eGFP expression levels in cells mock transfected or expressing mCherry, eGFP, mCherry and eGFP or an mCherry-eGFP fusion protein separated by 10 amino acids.

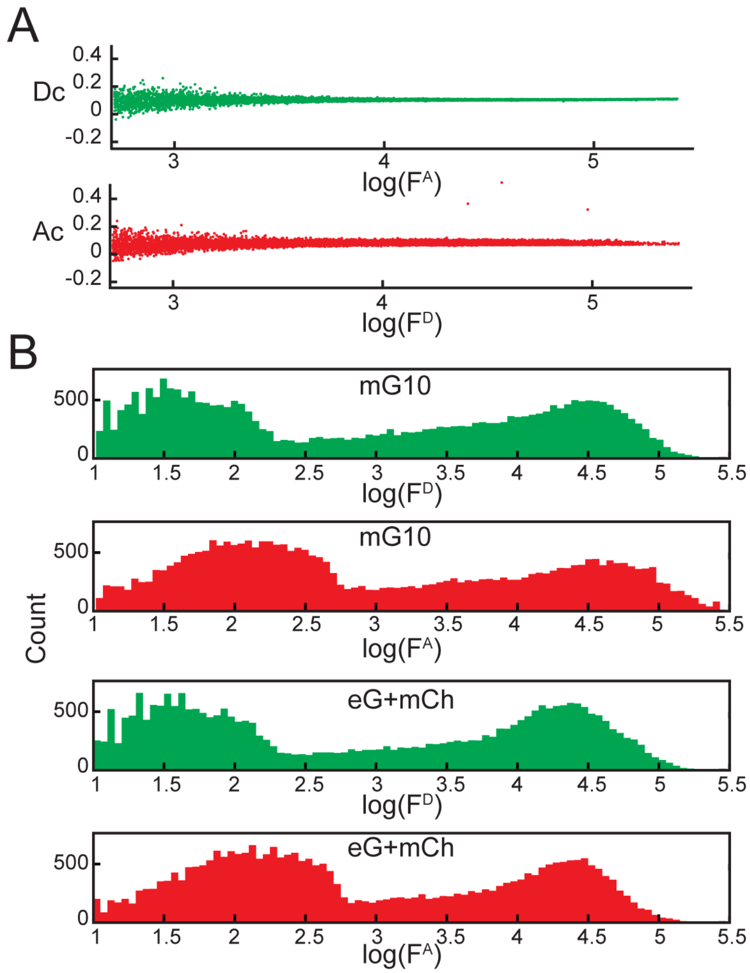


**Figure S4. Comparison of the cross-talk corrections and fluorescence distributions in cells expressing fluorescent proteins analyzed by multichannel flow cytometry**

A) Scatterplots of of the donor (Dc, top panel) and acceptor (Ac, bottom panel) cross-talk corrections used to adjust the fluorescence levels prior to calculation of FRET signal. The top panel uses fluorescence from cells expressing eGFP only while the lower panel is from cells expressing mCherry only. B) Histograms of mCherry and eGFP fluorescence levels in cells transfected with mG10 fusion protein (*upper pair*) or mCherry and eGFP (*lower pair*).

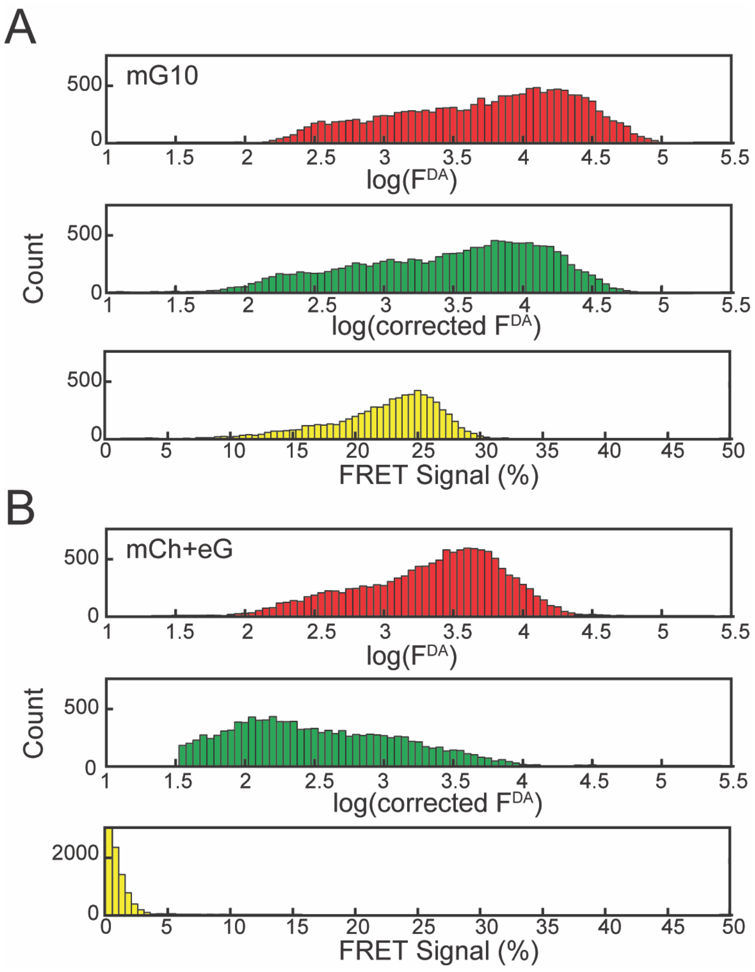


**Figure S5. Calculation of $N_{FRET}$ signals from flow cytometry intensities.**
Histograms of fluorescence intensities from HEK293T cells expressing mG10 (A) or mCherry and eGFP (B) in the FRET channel ($F^{DA}$, upper panels), corrected for cross-talk (corrected $F^{DA}$, *middle panels*) and converted to $N_{FRET}$ by normalization with donor and acceptor fluorescence (FRET Signal (%), *lower panels*).

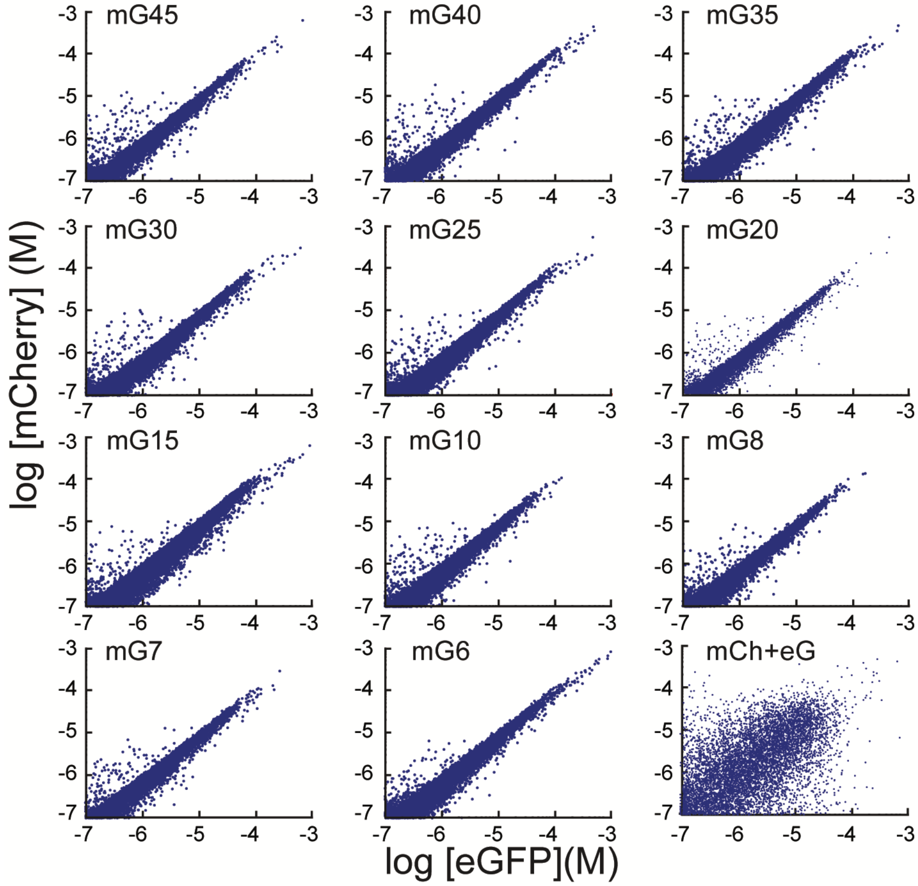


**Figure S6. Multichannel flow cytometry of HEK293T cells expressing mCherry-eGFP fusion proteins separated by linkers of different lengths.**
Histograms of multichannel fluorescence intensities converted to concentration from HEK293T cells expressing mG proteins.

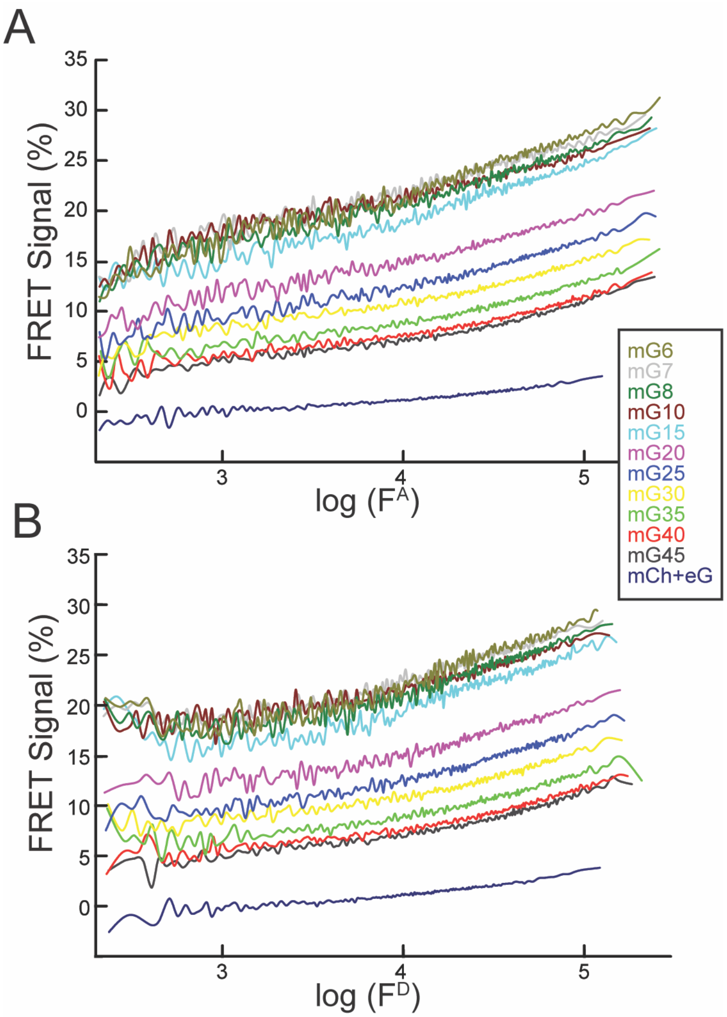


**Figure S7. FC-FRET signal dependence on donor and acceptor fluorescence intensities.** FRET signals from HEK293T cells expressing mG fusion proteins are plotted as a function of acceptor (A) and donor (B) fluorescence intensity. Intensity range was divided into bins of 100 cells and the mean $N_{FRET}$ signal is plotted as a moving average.

**Figure S8. Stochastic FRET between mCherry and eGFP.**

A. Calculation of the FRET efficiency as a function of acceptor concentration (assuming a 1:1 donor-acceptor ratio) arising from molecular collisions of freely diffusing eGFP and mCherry using the Lakowicz equation (*see* Methods). The various values of $R_0$ are indicated. B. Simulation including protein exclusion size. An example distribution of donor and acceptor molecules (100) randomly arranged in a sphere with a diameter of 100 nm. The distribution was generated including an exclusion zone for each fluorophore to simulate the volume of the protein core. Those pairs that met the distance criterion between the two fluorophores are connected (*lines*). C) For Monte Carlo simulation of stochastic FRET, random distributions as in (B) with different concentrations of donor-acceptor pairs were generated, the ensemble FRET efficiency calculated and then averaged to generate the relative FRET efficiency. The actual measured $N_{FRET}$ from an FC-FRET experiment is shown (*dotted line*).

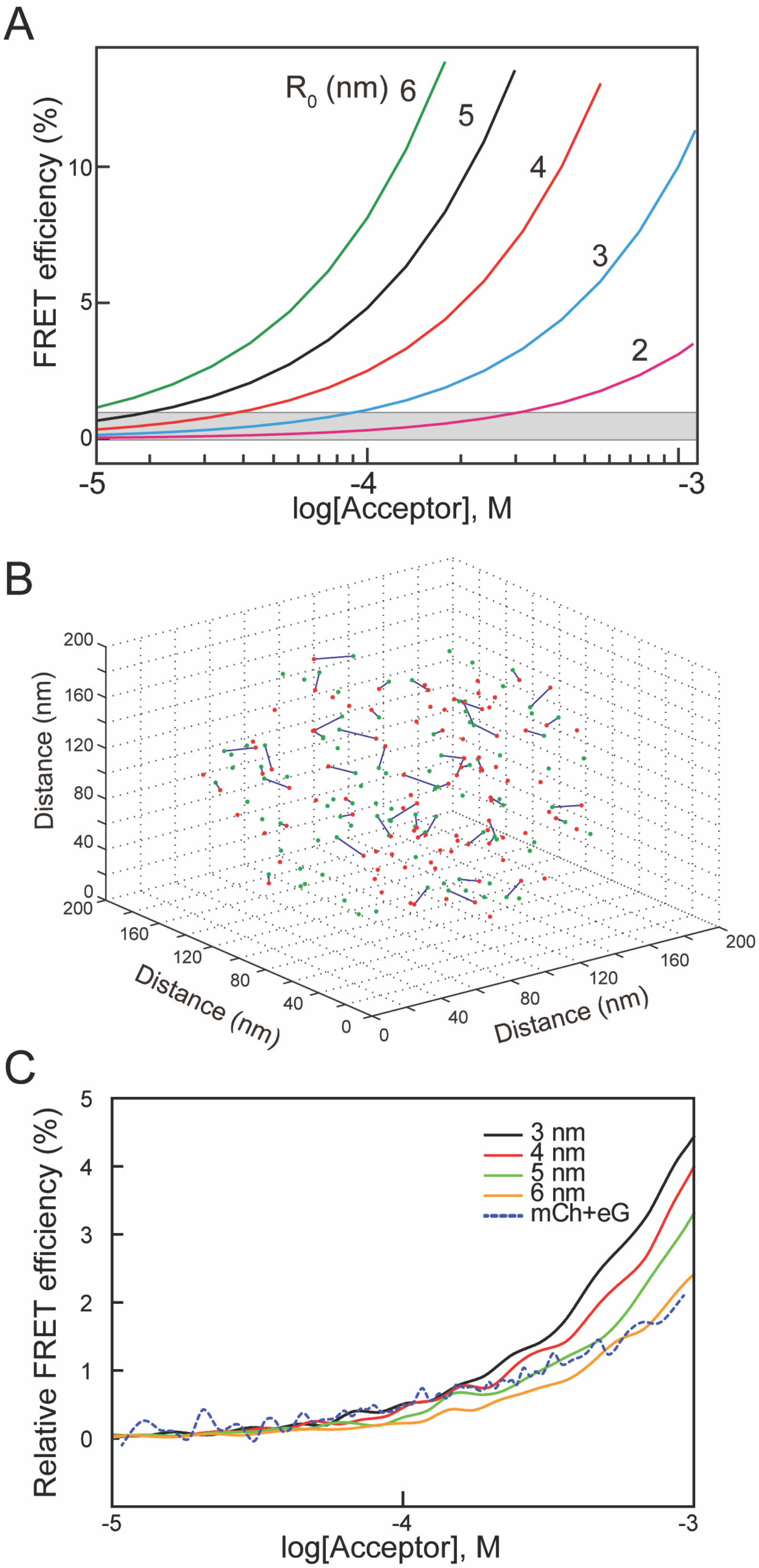

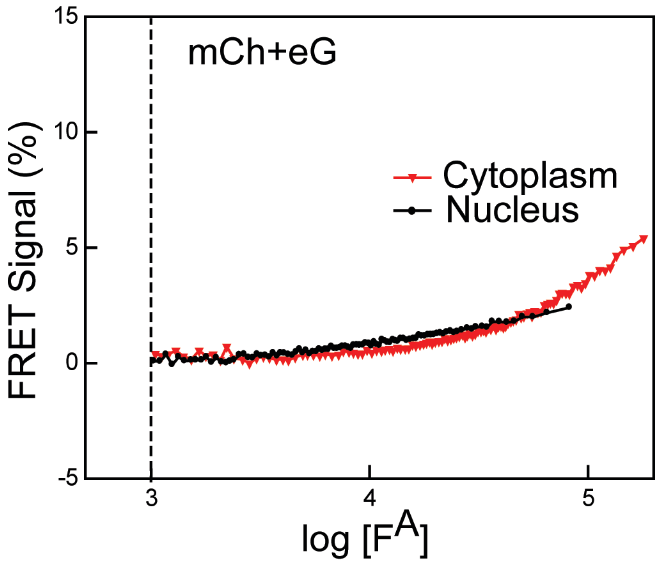


**Figure S9. Comparison of FC-FRET signals from mCherry and eGFP co-expressed in HEK293T cells.**

HEK293T cells were co-transfected with GFP and mCherry with (*black*) and without the nuclear localization signal (red). The $N_{FRET}$ signal was determined for each cell, the cells were sorted by acceptor intensity and divided into bins of 100 cells. The symbols are the mean for each bin and the line a moving average.

**Supplemental Table 1**
**Linkers for the mCherry-eGFP fusion proteins**

| Name | Length aa | Amino Acid Sequence |
|---|---|---|
| mG6 | 6 | $NH_2$-DPPVAT |
| mG7 | 7 | $NH_2$-DPVPVAT |
| mG8 | 8 | $NH_2$-DPAVPVAT |
| mG10 | 10 | $NH_2$-GGGLDPPVAT |
| mG15 | 15 | $NH_2$-GGGLDPPVATGGGGG |
| mG20 | 20 | $NH_2$-DPGA$(EAAAK)_2$AVPVAT |
| mG25 | 25 | $NH_2$-DPGA$(EAAAK)_3$AVPVAT |
| mG30 | 30 | $NH_2$-DPGA$(EAAAK)_4$AVPVAT |
| mG35 | 35 | $NH_2$-DPGA$(EAAAK)_5$AVPVAT |
| mG40 | 40 | $NH_2$-DPGA$(EAAAK)_6$AVPVAT |
| mG45 | 45 | $NH_2$-DPGA$(EAAAK)_7$AVPVAT |

**Supplemental Table 2**
**Linker sequences for the Crx and Nrl mCherry/eGFP fusion proteins**

| Construct | Amino Acid Sequence |
|---|---|
| **eNrl** | $NH_2$-MAPKKKRKVNRSKAEQKLISEEDLNSRPLE-**eGFP**-GGGGGNSSR(L)-**Nrl** |
| **eCrx** | $NH_2$-MAPKKKRKVNRSKAEQKLISEEDLNSRPLE-**eGFP**-GRWRWRPR-Crx |
| **Nrle** | $NH_2$-MAPKKKRKVNRSKAEQKLISEEDLNSS(R)-**Nrl**-GGGGG-**eGFP** |
| **Crxe** | $NH_2$-MAPKKKRKVNRSKAEQKLISEEDLNSSRPLE-**Crx**-GGGGG-**eGFP** |
| **mNrl** | $NH_2$-MAPKKKRKVNRSKAEQKLISEEDLNSGGGG-**mCherry**-GGGGGNSS(R)-**Nrl** |
| **mCrx** | $NH_2$-MAPKKKRKVNRSKAEQKLISEEDLNSRPLEG-**mCherry**-GGGGGGLE-**Crx** |
| **Nrlm** | $NH_2$-MAPKKKRKVNRSKAEQKLISEEDLNSS(R)-**Nrl**-GGGGG-**mCherry** |
| **Crxm** | $NH_2$-MAPKKKRKVNRSKAEQKLISEEDLNSRPLE-**Crx**-GTAGPGSGGGGG-**mCherry** |

The nuclear localization sequence is underlined. The NCBI accession numbers are for human Crx, NP_000545 and human Nrl, NP_001341697. For Nrl constructs, the initiator codon was replaced by the amino acid in parentheses. eGFP and mCherry cDNAs were from Clontech.
